# Neonatal Delivery of Exon Skipping Therapeutics Maximizes Brain Dystrophin Rescue in the *Mdx52* Mouse Model of Duchenne Muscular Dystrophy

**DOI:** 10.64898/2026.10.09.757591

**Authors:** Ophélie Vacca, Mathilde Doisy, Cathy Nagy, Amel Saoudi, Xaysongkhame Micky Phongsavanh, Olivier Le Coz, Cyrille Vaillend, Aurélie Goyenvalle

**Affiliations:** Université Paris-Saclay, UVSQ, Inserm, UMR 1357 IMPROVE, 78000 Versailles, France; Université Paris-Saclay, CNRS, Institut des Neurosciences Paris-Saclay, 91400 Saclay, France

**Keywords:** Exon skipping, Antisense Oligonucleotides (ASO), Adeno-Associated Virus (AAV), Central nervous system (CNS), Duchenne muscular dystrophy, Dystrophin, *Mdx52* Mouse Model, Intracerebroventricular Injection, Systemic Injection

## Abstract

Duchenne muscular dystrophy (DMD) is caused by mutations in the *DMD* gene, resulting in the loss of dystrophin expression. Beyond progressive muscle degeneration, nearly half of patients exhibit cognitive and behavioral impairments, highlighting the need for effective central nervous system (CNS) therapies. In this study, we investigated whether neonatal delivery could maximize brain dystrophin rescue in *mdx52* mice using exon 51 skipping strategies. We compared systemic and intracerebroventricular (ICV) delivery of an antisense oligonucleotide (ASO-51) and an AAV9-mediated U7snRNA approach (scAAV9-U7-Ex51), administered neonatally. Neonatal intravenous ASO-51 induced widespread but transient exon skipping, resulting in limited brain dystrophin restoration. ICV administration markedly increased CNS exon skipping but did not overcome the limited persistence of ASO, leading to progressive loss of exon skipping and dystrophin rescue. In contrast, neonatal ICV delivery of scAAV9-U7-Ex51 achieved robust, widespread, and sustained exon 51 skipping throughout the brain, including the hippocampus, cortex, and cerebellum, resulting in 20–50% restoration of Dp427 expression. These findings demonstrate that early CNS-directed delivery enhances brain dystrophin restoration and establish AAV-U7-mediated exon skipping as a promising strategy for addressing CNS comorbidities in DMD.

## INTRODUCTION

Nucleic acid-based therapeutics have recently shown significant promise for targeting the central nervous system (CNS), opening new avenues to address neurological manifestations of genetic disorders. Among these, antisense oligonucleotides (ASOs) and adeno-associated viral (AAV) vectors have emerged as prominent approaches, each offering distinct advantages in modulating gene expression in neural tissues. While ASOs enable direct and flexible modulation of RNA processing, their effects are often transient and may require repeated administration. In contrast, AAV-based approaches allow sustained expression of therapeutic sequences following a single administration. In particular, AAV9 has demonstrated transduction efficiencies comparable to other CNS-targeting serotypes when delivered via intracerebroventricular (ICV) injection^1^, supporting its suitability for direct CNS targeting. Complementing these approaches, U7snRNA-based systems enable the stable expression of antisense sequences from a U7 cassette under the control of its endogenous promoter, thereby combining the specificity of antisense-mediated splicing modulation with the durability of viral gene transfer^2,3^. In the context of Duchenne muscular dystrophy, AAV-U7 exon-skipping strategies have shown particularly promising results, supporting durable dystrophin restoration in different disease models and highlighting their potential for addressing both muscular and neurological manifestations of the disease^4,5^. Together, these approaches outline a versatile therapeutic framework for targeting RNA processing in the CNS.

Duchenne muscular dystrophy (DMD) is an X-linked disorder caused by mutations in the *DMD* gene that abolish functional dystrophin and drive progressive skeletal muscle degeneration and cardiomyopathy. Although traditionally viewed as a peripheral myopathy, DMD is now recognized as a multisystem disease with clinically meaningful CNS involvement. Approximately 40-50% of affected individuals exhibit neurocognitive and neurobehavioral comorbidities as lower IQ, executive dysfunction, attentional deficits, autism spectrum traits, anxiety, and emotional dysregulation, highlighting a major unmet need insufficiently addressed by current muscle-directed therapies^6–9^. Multiple brain dystrophins, including full-length Dp427, Dp140, and Dp71, are differentially expressed across neural cell types and regions, where they scaffold synaptic complexes and contribute to inhibitory/excitatory neurotransmission and circuit maturation^10^. Recent human and mouse atlases map dynamic, isoform-specific spatiotemporal patterns during development and adulthood, with Dp427 and Dp140 expression linked to neuronal populations critical for learning and emotional regulation^11,12^. Loss of these isoforms perturbs GABAA receptor clustering and synaptic organization, providing mechanistic substrates for DMD brain phenotypes^13^.

Exon-skipping ASOs aimed at restoring the open reading frame of *DMD* transcripts are clinically approved for subgroups of patients (e.g., targeting exon 51, 53, 45). While these drugs increase truncated dystrophin in skeletal muscle, effective and sustained brain rescue remains limited by delivery barriers, notably the blood-brain barrier (BBB), leaving CNS comorbidities largely untreated^14,15^.

Direct cerebrospinal fluid (CSF) administration (intracerebroventricular, ICV; intrathecal) can bypass the BBB and has shown broad CNS biodistribution for ASOs in rodents and large animals, with ICV favoring deeper brain penetration in comparative studies^16,17^. In adult *mdx52* mice, ICV delivery of tricyclo-DNA ASOs targeting exon 51 was previously shown to induce substantial exon skipping and partial Dp427 restoration throughout the brain, demonstrating the feasibility of CNS-targeted exon-skipping approaches in DMD^18^. Nevertheless, protein restoration remained heterogeneous and incomplete, raising the possibility that intervention during earlier developmental stages could further enhance therapeutic efficacy.

Vectorized exon skipping via AAV-U7 small nuclear RNA (U7snRNA) offers sustained splice redirection and has achieved widespread, durable dystrophin restoration in muscle across DMD models^4,5^. In *mdx52* mice, a model lacking Dp427 and Dp140 that recapitulates key DMD brain phenotypes, adult CNS delivery of AAV9-U7 for exon 51 drives exon skipping but yields variable protein rescue and limited behavioral correction, suggesting that earlier intervention may be required to capitalize on developmental windows for neural circuit formation^18,19^.

Biological and developmental considerations suggest that therapeutic timing may be decisive for CNS benefit. BBB properties emerge early in development and mature postnatally through neurovascular unit interactions; although the BBB is functional in utero, its architecture and transport systems evolve over the perinatal period, with implications for biodistribution of macromolecules and viral vectors^20,21^.Here, we aimed to maximize exon-skipping efficacy in the CNS of *mdx52* mice by achieving widespread dystrophin restoration as the earliest possible postnatal stage, thereby overcoming the limitations associated with later interventions. To this end, we evaluated complementary exon-skipping approaches and delivery routes in neonatal mice. Specifically, we compared intravenous and ICV administration of an ASO and an AAV9-U7snRNA vector both targeting *Dmd* exon 51 in *mdx52* mice, and quantified exon skipping and Dp427 restoration across major brain regions, including the hippocampus, cortex, and cerebellum, as well as in peripheral tissues. By systematically assessing the extent of dystrophin restoration, its biodistribution and long-term persistence, our aim was to identify the most effective strategy for early CNS dystrophin rescue and provide the evidence needed to guide the development of CNS-targeted therapies for DMD neurobehavioral manifestations.

## RESULTS

### ASO-51 intravenous injection into the temporal vein of *mdx52* neonates

We first performed a single intravenous administration of ASO-51 at postnatal day 2 (P2) in *mdx52* mice and analysed the biodistribution, and levels of exon skipping and dystrophin restoration at different time points after injections (week 4, 7 and 14 post injection) (Figure 1A). Systemic injection of ASO resulted in a widespread and time-dependent distribution of the oligonucleotide across the CNS and peripheral tissues in *mdx52* mice (Figure 1B). Within the CNS, ASO accumulation was detectable in the hippocampus (HIP), cortex (CX), cerebellum (CBL) and spinal cord (SC) as early as week 4 (W4), with relatively stable levels observed at W7 and W14 despite high inter-individual variability (Figure 1B and S1A). In contrast, peripheral tissues which displayed higher ASO concentrations overall, particularly in the diaphragm (DIA) and heart, were characterized by a strong decline between W4 and later time points (Figure 1C) (time effect P value < 0,0001 analysed by two-way ANOVA).

**Figure 1:**
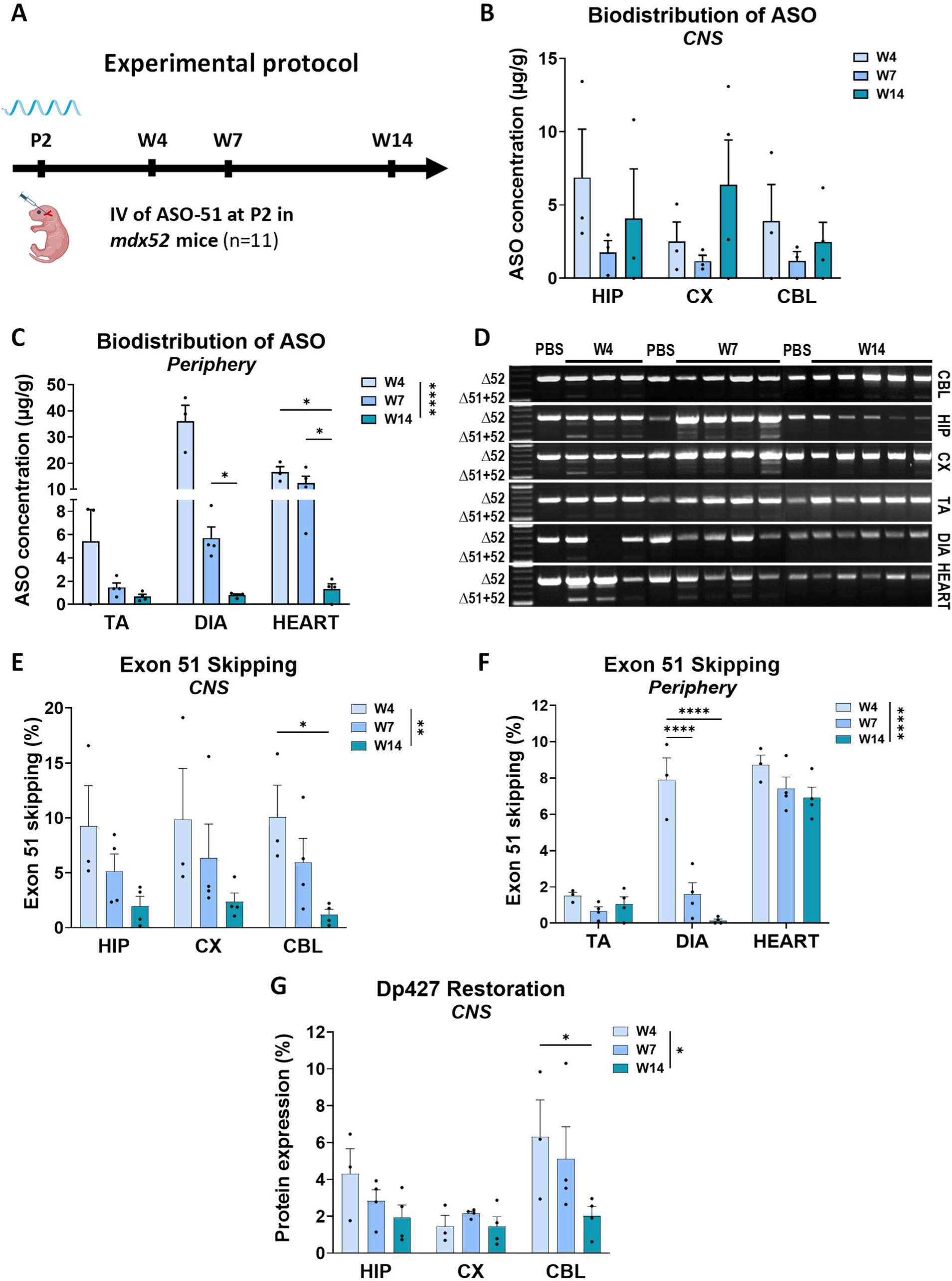
Systemic injection of ASO-51 in *mdx52* at P2 and longitudinal molecular outcomes. (A) Experimental design. *Mdx52* mice received a single intravenous injection of ASO-51 (1.2 mg) into the superficial temporal vein at postnatal day 2 (P2). Tissues were collected 4, 7, or 14 weeks after injection (W4, W7, W14). (B-C) ASO-51 biodistribution across central and peripheral tissues at W4, W7 and W14. (D) Representative RT-PCR gels showing the skipped Δ51+52 product and the unskipped Δ52 amplicon in central nervous system CNS) and peripheral tissues, illustrating exon 51 skipping efficiency at each time point. (E) Quantification of exon 51 skipping in CNS regions including hippocampus (HIP), cortex (CX), cerebellum (CBL) by RT-qPCR at W4, W7, and W14. (F) Quantification of exon 51 skipping in peripheral tissues (tibialis anterior [TA], diaphragm [DIA], heart) by RT-qPCR at the same time points. (G) Dp427 protein restoration in CNS regions quantified by western blot and expressed as a percentage of wild-type level. Data are presented as mean ± SEM with individual points shown. Statistical analysis was performed using two-way ANOVA followed by Tukey’s multiple comparisons test. *p < 0.05, **p < 0.01, ****p < 0.001. The sample sizes for each time point were as follows: W4 (n = 3), W7 (n = 4), and W14 (n = 4).

RT-PCR and RT-qPCR analyses confirmed successful exon 51 skipping in both CNS and peripheral tissues (Figure 1D-F). In the CNS, exon skipping levels were modest but detectable in all regions examined, with the highest levels observed at W4 (mean ∼10%) and a progressive decrease at W7 and W14 (Figure 1E and S1B). In peripheral tissues, exon 51 skipping was variable across muscles (Figure 1F). The diaphragm showed the strongest response, reaching ∼8% skipping at W4, which significantly decreased at later time points. The heart also exhibited robust exon skipping (∼6-8%) which remain stable across all time points, whereas tibialis anterior (TA) muscle showed comparatively very low levels.

Consistent with exon skipping levels, Dp427 protein restoration was detected in CNS tissues (Figure 1G and S2A). Protein expression was the highest at W4, particularly in the cerebellum (∼6-7% of wild-type levels), and declined over time, while remaining detectable at W14. The hippocampus showed moderate restoration (∼3-5%), and the cortex exhibited the lowest levels overall.

Collectively, these results demonstrate that early systemic delivery of ASO-51 enables broad biodistribution and detectable exon skipping and dystrophin restoration in the CNS, albeit at relatively low levels and with a time-dependent decline.

### scAAV9-U7-Ex51 intravenous injection into the temporal vein of *mdx52* neonates

To circumvent the transient effect of ASO, we next evaluated a vectorized approach using the AAV-U7snRNA system. *Mdx52* mice were injected intravenously at P2 with the previously described scAAV9-U7-Ex51^21^ and we analysed vector distribution, and levels of exon skipping and dystrophin restoration across both CNS and peripheral tissues at different time points after injections (week 4, 7 and 14 post injection) (Figure 2A). Quantification of viral genomes revealed low CNS targeting, with the highest levels observed in the hippocampus (HIP), followed by the cortex (CX), cerebellum (CBL) and the spinal cord (Figure 2B and S1C). Although viral genomes were detectable up to W14, CNS levels remained insufficient compared with peripheral tissues, indicating poor brain targeting following systemic neonatal administration. In peripheral tissues, vector distribution was markedly higher overall compared to the CNS (Figure 2C). The heart exhibited the highest viral genome copy numbers per cell at all time points, reaching approximately 11 vg/cell at W14, whereas the diaphragm (DIA) and tibialis anterior (TA) showed lower but detectable levels.

**Figure 2:**
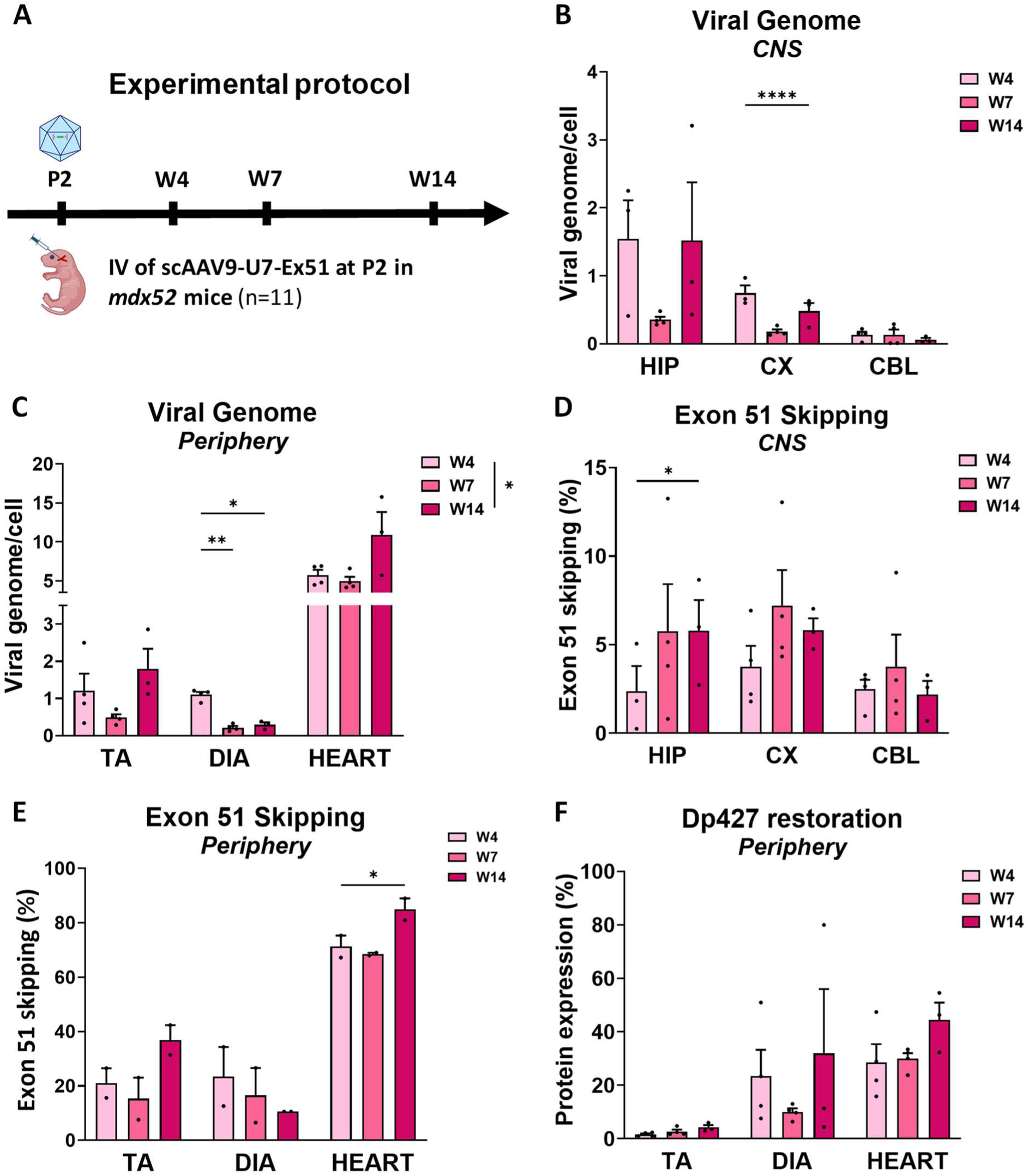
Systemic injection of scAAV9-U7-Ex51 in *mdx52* at P2 and longitudinal molecular outcomes. (A) Experimental timeline. *Mdx52* mice received a single intravenous injection of 2E+12 vg of scAAV9-U7-Ex51 into the superficial temporal vein at postnatal day 2 (P2). Tissues were collected at 4, 7, and/or 14 weeks post-injection. (B-C) Viral genome quantification by qPCR in CNS regions (B) including hippocampus (HIP), cortex (CX), and cerebellum (CBL), and in peripheral tissues (C) including tibialis anterior (TA), diaphragm (DIA), and heart, at the indicated time point(s). (D-E) Exon 51 skipping quantification by RT-qPCR in (D) CNS regions including hippocampus (HIP), cortex (CX), and cerebellum (CBL), and in (E) peripheral tissues including tibialis anterior (TA), diaphragm (DIA), and heart. (F) Restoration of dystrophin Dp427 protein expression in peripheral tissues, quantified by Jess simple western and expressed as a percentage of wild-type levels. Data are presented as mean ± SEM with individual data points shown. Statistical analysis was performed using a two-way ANOVA followed by Tukey’s multiple comparisons test. *p < 0.05, **p < 0.01, ****p < 0.0001. The sample sizes for each time point were as follows: W4 (n = 4), W7 (n = 4), and W14 (n = 3).

Consistent with vector biodistribution, exon 51 skipping was detected at relatively low levels in all CNS regions analyzed (Figure 2D and S1D), ranging from approximately 4% to 8%, with the cortex showing the highest values across time points. In peripheral tissues, exon skipping levels were substantially higher than in the CNS (Figure 2E). The heart demonstrated the most pronounced effect, reaching up to ∼80-90% exon skipping at W14. The diaphragm exhibited intermediate levels (∼15-25%), while the tibialis anterior showed lower but consistent skipping (∼20-35%), with some variability across time points.

These exon skipping levels translated into robust dystrophin Dp427 restoration in peripheral tissues (Figure 2F and S2B), with the highest protein expression detected in the heart, reaching approximately 40-50% of wild-type levels at W14 and more moderate restoration in the diaphragm and tibialis anterior (∼30% and ∼4% respectively). However, despite detectable exon 51 skipping in all CNS regions, no Dp427 restoration could be detected by western blot analysis (Figure S2B). This observation is consistent with previous studies using systemic AAV9-U7-mediated exon skipping in *mdx52* mice, where low levels of brain exon skipping were similarly insufficient to generate detectable dystrophin protein restoration despite persistent splice correction^5^. These findings suggest that the levels of exon skipping achieved in the CNS following systemic neonatal administration remain below the threshold required for measurable dystrophin recovery and indicate that early systemic administration of scAAV9-U7-Ex51 is more appropriate for targeting peripheral tissues, particularly the heart, than CNS regions.

### Bilateral ICV injection of ASO-51 in *mdx52* neonates

Given the primary objective of this study to achieve effective and widespread exon skipping and dystrophin restoration in the brain of *mdx52* mice, we next evaluated direct ICV administration in *mdx52* neonates. We first performed bilateral ICV injection of ASO-51 in *mdx52* at P1 and like above, analysed the biodistribution, and levels of exon skipping and dystrophin restoration at different time points after injections (week 4, 7 and 14 post injection) (Figure 3A). Direct ICV administration of ASO-51 resulted in efficient and widespread delivery throughout the CNS of *mdx52* mice. High levels of ASO were detected in all analysed regions at W4, with the cortex (CX) exhibiting the highest concentrations, reaching approximately 40 µg/g at W4, followed by hippocampus (HIP) showing ∼20 µg/g and cerebellum (CBL) displaying lower but detectable concentrations around 5-10 µg/g (Figure 3B). Yet, the levels were significantly lower at W14 suggesting progressive clearance of the ASO (time effect p-value < 0.0001 analysed by two-way ANOVA). Comparable ASO accumulation was also observed in the spinal cord (SC), supporting widespread distribution across the neuraxis following ICV delivery (Figure S1E). In contrast with CNS regions, peripheral exposure was minimal following ICV delivery of ASO with low ASO levels detected in diaphragm (DIA) and heart, and negligible amount the tibialis anterior (TA), indicating limited systemic distribution (Figure S3).

**Figure 3.**
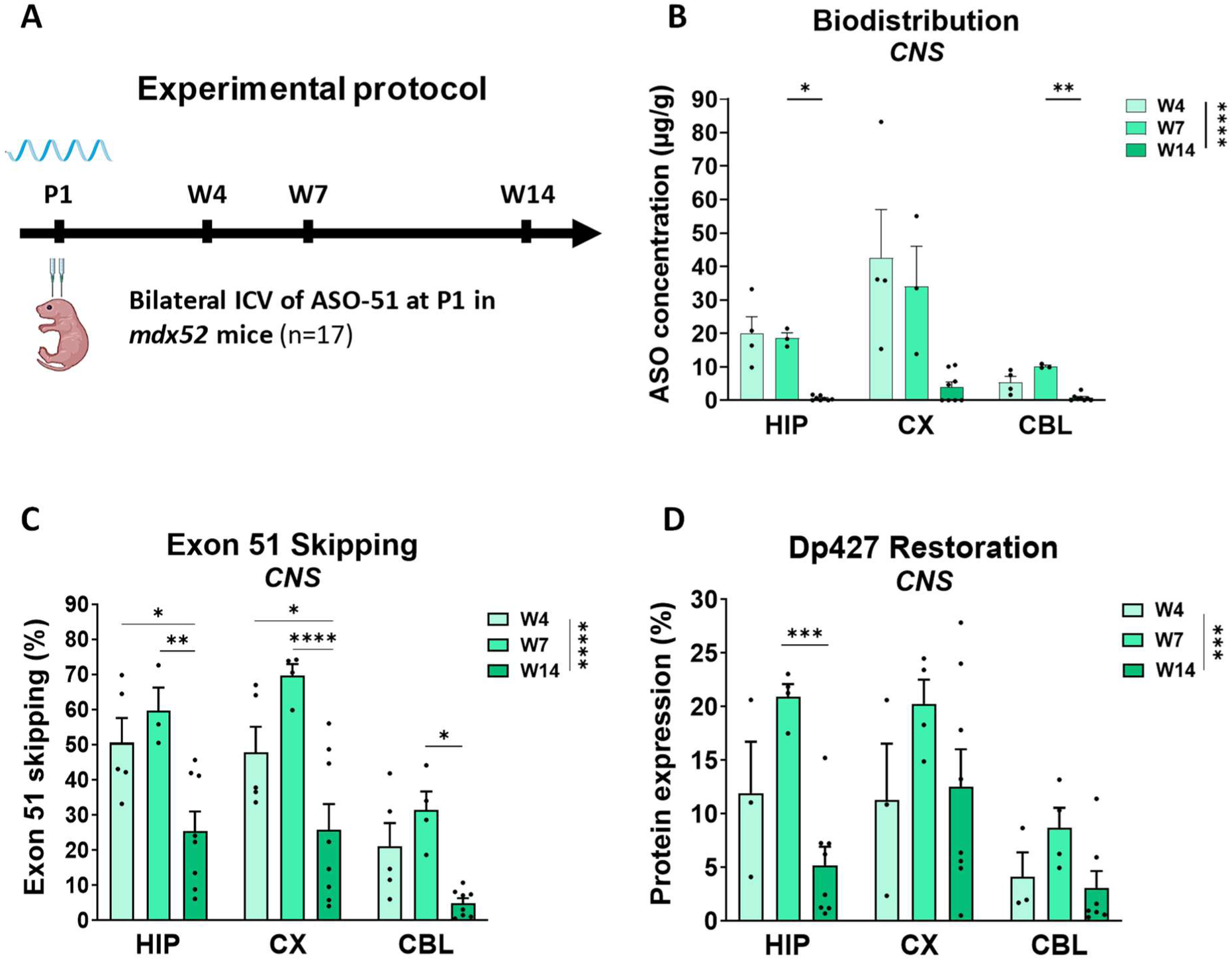
Bilateral intracerebroventricular (ICV) injection of ASO-Ex51 in *mdx52* at P1. (A) Experimental timeline. *Mdx52* mice received a single bilateral intracerebroventricular (ICV) injection of ASO-Ex51 (160 µg) at postnatal day 1 (P1). Tissues were collected at 4, 7 or 14 weeks post-injection (W4, W7, W14). (B) ASO-51 biodistribution in CNS regions, including hippocampus (HIP), cortex (CX), and cerebellum at W4, W7 and W14. (C) Exon 51 skipping quantification by RT-qPCR in hippocampus (HIP), cortex (CX), and cerebellum (CBL), at W4, W7, and W14. (D) Restoration of dystrophin Dp427 protein expression in CNS regions, quantified by western blot and expressed as a percentage of wild-type levels. Data are presented as mean ± SEM with individual data points shown. Statistical analysis was performed using a two-way ANOVA followed by Tukey’s multiple comparisons test. *p < 0.05, **p < 0.01, ***p < 0.001, ****p < 0.0001. The sample sizes for each time point were as follows: W4 (n = 5), W7 (n = 4), and W14 (n = 8).

Consistent with the CNS biodistribution, exon 51 skipping levels were markedly elevated in all brain regions (Figure 3C). The cortex demonstrated the highest skipping levels, reaching ∼70-80% at W7 and declining to ∼20-30% by W14. The hippocampus also showed robust exon skipping ∼50-60% at W7, declining to ∼20-30% by W14, while the cerebellum exhibited lower but substantial levels ∼25-40%, declining to ∼5-10% by W14. In the spinal cord (SC), exon 51 skipping reached approximately ∼60% at W7, decreasing to ∼20-25% at W14, consistent with the transient profile observed in brain regions (Figure S1F).

Importantly, this robust exon skipping in the CNS translated into significant Dp427 protein restoration (Figure 3D and S2C). Protein expression peaked at W7 in all regions, reaching approximately 20-25% of wild-type levels in the hippocampus and cortex, and ∼8-13% in the cerebellum. However, Dp427 protein levels markedly declined by W14, particularly in the hippocampus, consistent with ASO biodistribution and exon skipping levels. Collectively, these findings demonstrate that early ICV administration of ASO-51 enables robust CNS rescue, with high levels of dystrophin restoration at W7. However, the marked decline observed by W14 indicates that this effect is not durable, and therefore may be insufficient to support long-term correction of cognitive deficits in *mdx52* mice.

### Bilateral ICV injection of scAAV9-U7-Ex51 in *mdx52* neonates

In order to achieve persistent exon skipping and dystrophin restoration, we next performed direct ICV delivery of scAAV9-U7-Ex51 in *mdx52* mice at P1 which enabled efficient and sustained transduction of the CNS in *mdx52* mice (Figure 4A). Viral genome quantification confirmed robust vector distribution throughout the CNS (Figure 4B and S1G). The highest levels were observed in the cortex (CX), reaching approximately 20-25 vg copies per cell at W7, followed by the hippocampus (HIP), which displayed stable levels (∼10-15 vg/cell) across time points. Intermediate viral genome levels were also detected in the spinal cord (SC), with approximately ∼4-6 vg/cell across time points, supporting CNS-wide vector dissemination. In contrast, the cerebellum (CBL) showed lower vector presence, with approximately 0.4 vg copies per cells.

**Figure 4.**
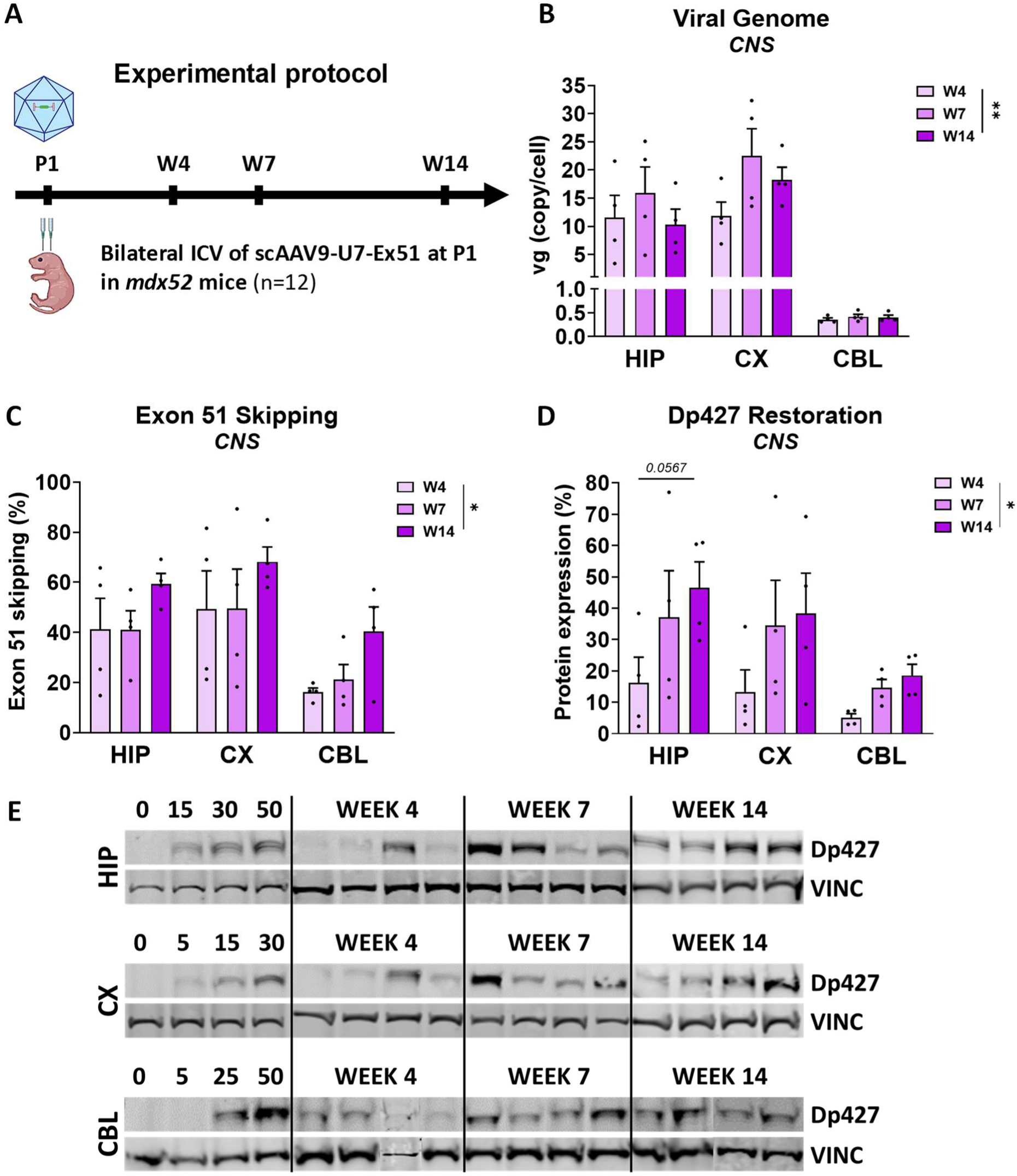
Bilateral intracerebroventricular (ICV) injection of scAAV9-U7-Ex51 in *mdx52* at P1. (A) Experimental timeline. *Mdx52* mice received a single bilateral intracerebroventricular (ICV) injection of scAAV9-U7-Ex51 (7E+11 vg) at postnatal day 1 (P1). Tissues were collected at 4, 7, or 14 weeks post-injection (W4, W7, W14). (B) Viral genome quantification by qPCR in CNS regions including hippocampus (HIP), cortex (CX), and cerebellum (CBL) at W4, W7, and W14. (C) Exon 51 skipping quantification by RT-qPCR in hippocampus (HIP), cortex (CX), and cerebellum (CBL) at W4, W7, and W14. (D) Quantification of dystrophin Dp427 protein expression in CNS regions, by western blot and expressed as a percentage of wild-type levels. (E) Representative gels of western blot detecting Dp427 expression and vinculin as a loading control in hippocampus (HIP), cortex (CX), and cerebellum (CBL) at W4, W7, and W14. Data are presented as mean ± SEM with individual data points shown. Statistical analysis was performed using a two-way ANOVA followed by Tukey’s multiple comparisons test. *p < 0.05, **p < 0.01. The sample sizes for each time point were as follows: W4 (n = 4), W7 (n = 4), and W14 (n = 4).

Consistent with these biodistribution profiles, exon 51 skipping was efficiently induced in all CNS regions (Figure 4C and S1H). The cortex exhibited the highest skipping levels, reaching ∼60-70% at W14, followed by the hippocampus showing ∼40-60% across time points and the cerebellum with more moderate exon skipping levels (∼15-40%). Interestingly, levels of exon skipping appeared to increase with time (time effect p-value < 0.0001 analysed by two-way ANOVA), likely due to the continuous expression of the U7-Ex51 in transduced cells. In the spinal cord (SC), exon 51 skipping levels followed the same pattern, reaching approximately 20% at W14 (Figure S1H).

In peripheral tissues, exon skipping remained very limited following ICV of scAAV9 delivery, with less than 2% detected in tibialis anterior (TA) and diaphragm (DIA) at W14, except in the heart where significant exon 51 skipping was detected (∼30%) (Figure S4).

Importantly, Western blot analysis confirmed that the robust exon skipping observed in the CNS translated into substantial Dp427 protein restoration (Figure 4D-E). Protein expression increased progressively over time, particularly in the hippocampus and cortex, reaching ∼40-50% and ∼35-40% of wild-type levels, respectively, by W14. The cerebellum showed lower but significant dystrophin restoration (∼15-20%), consistent with lower viral genome levels and exon skipping levels in this region.

Overall, these results demonstrate that ICV delivery of scAAV9-U7-Ex51 at early postnatal stages results in strong and durable CNS transduction, leading to high levels of exon skipping and substantial dystrophin restoration.

## DISCUSSION

In this study, we demonstrate that both the therapeutic modality and the route of administration are critical determinants of achieving widespread, durable exon 51 skipping and meaningful Dp427 restoration across multiple brain regions in *mdx52* mice. It should be noted that exon 51 skipping restores only the full-length Dp427 isoform, as exon 51 contains the translation initiation codon for Dp140^22^, which is consequently lost during the exon-skipping process. Accordingly, restoration of the Dp140 isoform is not expected with this therapeutic strategy and was therefore not evaluated in the present study.

Neonatal intravenous administration of ASO-51 resulted in only low levels of exon 51 skipping and minimal Dp427 restoration in the CNS, whereas direct ICV administration of the same ASO markedly increased exon skipping and dystrophin restoration throughout the brain. However, despite the improved CNS delivery, ICV administration did not overcome the limited tissue persistence of ASOs, leading to a progressive decline in exon skipping and Dp427 restoration over time. These findings illustrate the challenges of achieving sustained CNS correction with ASO therapy in the context of our long-term objective of investigating the impact of dystrophin restoration on neurobehavioral phenotypes.

The transient nature of ASO-mediated exon skipping is well recognized and can be addressed clinically through repeated intrathecal administration, as routinely performed for antisense therapies such as nusinersen in patients with spinal muscular atrophy. In the *mdx52* mouse model, however, maintaining therapeutic levels throughout the period of behavioral assessment would require an additional ICV administration by stereotaxic surgery at approximately 12–14 weeks of age. Such an intervention would represent a major confounding factor for subsequent neurobehavioral analyses. Consequently, our objective was to establish a therapeutic strategy capable of providing sustained brain dystrophin restoration following a single neonatal intervention.

In contrast to ASO treatment, the stable expression provided by scAAV9-U7-Ex51 following bilateral neonatal ICV administration enabled robust, widespread, and long-lasting exon 51 skipping, resulting in sustained Dp427 restoration throughout the brain (up to ∼20–50%). Importantly, similar trends were observed in the spinal cord, demonstrating efficient targeting across the neuraxis. By comparison, although systemic neonatal delivery of scAAV9-U7-Ex51 also produced durable exon skipping and dystrophin restoration, its effects were largely restricted to peripheral tissues, with dystrophin restoration ranging from approximately 20% in skeletal muscles to 80% in the heart. The absence of detectable dystrophin restoration in the CNS following systemic AAV delivery further underscores the limitations of peripheral administration for targeting brain pathology, at least when using the AAV9 serotype. These findings indicate that, despite efficient and durable exon skipping in peripheral tissues, particularly in heart, systemic neonatal administration of scAAV9-U7-Ex51 does not achieve the primary objective of the present study, namely robust dystrophin restoration throughout the brain. While the substantial cardiac rescue may have important therapeutic value in the context of DMD, it remains outside the scope of the CNS-directed strategy investigated here, which aims to address the neurological manifestations associated with dystrophin deficiency.

Compared with our previous study investigating scAAV9-U7-Ex51 in adult *mdx52* mice, neonatal delivery produced higher dystrophin restoration and markedly reduced inter-animal variability^23^. In adult animals, bilateral ICV administration of scAAV9-U7-Ex51 resulted in highly heterogeneous exon skipping and Dp427 restoration, with mean protein levels of only 6-12% across brain regions and no detectable behavioral benefit despite successful molecular correction^23^. The more robust and homogeneous rescue achieved in neonates therefore highlights the importance of therapeutic timing for efficient CNS targeting. These results reinforce the concept that developmental timing is a critical determinant of CNS efficacy for nucleic acid–based therapeutics. BBB barrier formation begins early during development but continues to mature postnatally; concomitantly, CSF dynamics, ventricular volume, and extracellular matrix composition in the neonatal brain are likely to facilitate the dispersion and uptake of vectors and oligonucleotides compared to later stages^21^. Together with heightened synaptogenesis and network plasticity, these factors define a window of opportunity during which therapeutic interventions may achieve maximal functional impact. In this context, early restoration of dystrophin may promote the re-establishment of synaptic architecture and stabilization of inhibitory/excitatory balance. These interpretations are consistent with contemporary models of BBB development and CNS mechanobiology as well as with studies demonstrating improved CNS delivery following early ICV administration^21,24^.

While neonatal ICV delivery of scAAV9-U7-Ex51 led to robust and long-lasting protein rescue, regional differences in Dp427 restoration were observed across the hippocampus, cortex, and cerebellum, likely reflecting a combination of CSF flow dynamics and region-specific cellular tropism. Interestingly, the highest levels of Dp427 restoration were observed in the hippocampus and cortex, mirroring the regional distribution of full-length dystrophin isoforms reported in recent human and mouse brain atlases^11,12^. This concordance suggests that neonatal ICV delivery of scAAV9-U7-Ex51 not only achieves efficient transduction of the CNS but also restores dystrophin preferentially in brain regions that normally exhibit the highest endogenous expression of Dp427. Such a distribution may represent an important advantage from a therapeutic perspective, as it targets neural circuits in which dystrophin is believed to play major roles in synaptic function and cognitive processing. In contrast, dystrophin restoration was comparatively lower in the cerebellum despite the known abundance and functional importance of Dp427 in this structure. This observation may be relevant for future behavioral studies, as cerebellar dysfunction has been implicated in several cognitive manifestations of DMD, including deficits in working memory, motor learning, and higher-order cognitive processing^25,26^. Consequently, incomplete rescue within this structure could potentially limit the extent of functional recovery despite robust restoration in forebrain regions.

Nevertheless, previous studies have shown that relatively modest levels of brain dystrophin restoration can be sufficient to normalize synaptic function and improve behavioral phenotypes in dystrophin-deficient mice, suggesting a nonlinear relationship between dystrophin abundance and functional outcome^27^. Indeed, even partial Dp427 re-expression (∼10-30% of wild-type levels) has been shown to normalize synaptic function and behavioral phenotypes in *mdx* models, suggesting a nonlinear relationship between dystrophin abundance and functional outcome^27^. The levels achieved in this study (∼20–50%) exceed these proposed thresholds, supporting their potential biological relevance for circuit stabilization in *mdx52*, which additionally lacks Dp140. Future studies should incorporate sensitive pharmacodynamic readouts, such as GABA_A receptor subunit localization and synaptic clustering, to better define the functional consequences of CNS dystrophin restoration^10,13^.

Importantly, these findings are complemented by a recent spatial transcriptomic analysis of the same brain samples, which provides additional mechanistic insight into the cellular and regional determinants of therapeutic efficacy^28^. Specifically, exon skipping events were found to occur predominantly in neuronal populations, with heterogeneous distribution across brain regions, including lower efficiency in hippocampal areas despite high baseline expression of full-length isoforms. Moreover, only a small subset of cells exhibited exon skipping, yet these cells showed marked restoration of dystrophin expression, suggesting that a limited population of rescued neurons may disproportionately contribute to overall protein recovery. Spatial mapping further revealed distinct distributions of full-length and shorter dystrophin isoforms, indicating that therapeutic efficacy may be influenced by baseline isoform expression patterns across cell types. In addition, treatment-specific immune responses were identified, with ASO administration associated with microglial activation, whereas no comparable response was observed following AAV-U7 delivery^28^. Together, these complementary datasets provide a more comprehensive understanding of the mechanisms underlying CNS-targeted exon-skipping therapy and establish a valuable framework for interpreting therapeutic efficacy at both the tissue and cellular levels.

Although neonatal ICV delivery of scAAV9-U7-Ex51 achieved predominantly CNS-directed transduction, detectable exon skipping was also observed in peripheral tissues, particularly in the heart (∼30%), whereas only minimal effects were detected in skeletal muscles such as the tibialis anterior and diaphragm. Although not the primary objective of this study, these findings are consistent with known CSF-systemic exchange^16^ and with reports of broader AAV9 biodistribution following neonatal administration^29,30^. While this suggests the possibility of combined CNS and peripheral benefit after direct CNS administration, careful dose optimization would be required to balance efficacy and safety in a translational perspective. In particular, early-life CNS-directed AAV delivery necessitates monitoring for potential neuroinflammatory responses, dorsal root ganglion toxicity, and off-target distribution^31^. Encouragingly, CSF-directed AAV administration has shown acceptable tolerability in multiple preclinical and early clinical studies^30,32–35^, and may reduce systemic exposure compared to systemic approaches.

While AAV9 was selected for its well-characterized CNS tropism and extensive preclinical use, the impact of capsid selection on exon-skipping efficiency remains an important area of investigation. Comparative studies following CSF administration have suggested broadly similar transduction profiles between AAV9 and engineered variants such as PHP.eB^1,36^, although these comparisons remain limited, particularly in neonatal settings. Given the expanding repertoire of CNS-targeted AAV capsids, future studies should assess whether alternative vectors can further enhance the extent, distribution, and durability of therapeutic effects in the developing brain.

Taken together, our comparative analysis of systemic versus intracerebroventricular delivery and ASO versus AAV-U7 therapeutic platforms demonstrates that both the route of administration and the therapeutic modality are critical determinants of efficient and sustained CNS dystrophin restoration. Our findings establish neonatal ICV delivery of AAV-U7 as the most effective strategy for achieving widespread, durable exon skipping and Dp427 restoration throughout the brain and spinal cord, providing a robust experimental platform for studying the contribution of CNS dystrophin deficiency to the neurological manifestations of Duchenne muscular dystrophy.

The primary objective of the present study was not to evaluate the behavioral consequences of CNS dystrophin restoration or to establish a clinically translatable delivery strategy, but rather to identify the conditions that maximize early, widespread, and long-lasting dystrophin rescue throughout the developing CNS. Establishing these optimal conditions was an essential prerequisite for future studies investigating whether sustained brain dystrophin restoration translates into improvements in the cognitive and behavioral abnormalities associated with DMD. The robust and durable CNS rescue achieved here now provides the experimental framework required to directly address these important questions and to guide the development of future CNS-targeted exon-skipping therapies.

## MATERIALS AND METHODS

### Animals and antisense oligonucleotides

The *mdx52* mouse model of X-linked muscular dystrophy, characterized by a deletion of exon 52 in the DMD gene, was developed by Dr. Katsuki Motoya’s team. This model was created by replacing exon 52 with a neomycin resistance gene, resulting in the absence of Dp427, Dp260, and Dp140 dystrophins, while maintaining the expression of Dp116 in peripheral nerves and Dp71 in the brain and retina. The strain was backcrossed with C57BL/6J mice for over eight generations to ensure genetic stability. This mouse line was generously shared by Prof. Toshikuni Sasaoka (Department of Comparative & Experimental Medicine/Brain Research Institute, Niigata University, Japan). Breeding pairs were provided to our laboratory by Dr. Jun Tanihata and Dr. Shin’ichi Takeda (National Center of Neurology and Psychiatry, Tokyo, Japan). In our facility at UVSQ in Montigny-le-Bretonneux (France), heterozygous females were bred with *mdx52* males to produce *mdx52* males and their C57BL/6J wild-type (WT) littermates. Genotyping was performed using PCR analysis of tail DNA. All animal care and experimental protocols adhered to European regulations (CEE 86/609/EEC and EU Directive 2010/63/EU) as well as French national guidelines (87/848) and were approved by the Paris Center-South Ethics Committee (no. 47).

A previously described tricyclo-DNA (tcDNA) antisense oligonucleotide (ASO-51) directed against the exon 51 exonic splicing enhancer of dystrophin pre-mRNA (position +48 to +62) was used throughout this study^18^. The compound was produced by SQY Therapeutics (Montigny-le-Bretonneux, France). The tcDNA backbone was fully phosphodiester and functionalized at its 5′ end with a palmitic acid group via a C6-amino spacer and a phosphorothioate bond, following the previously reported chemistry.^37^

### Construction of U7snRNA Vectors and AAV Production

U7snRNA constructs targeting the murine *Dmd* exon 51 were engineered based on the previously described U7smOPT-SD23/BP22 backbone^38^ (modified murine U7snRNA gene). The antisense sequence originally targeting human exon 51 was adapted to the murine sequence by site-directed mutagenesis, introducing two specific nucleotide changes. The U7-ex51 long1 construct contains a single 45-mer antisense sequence complementary to positions +59 to +103 of murine exon 51^23^. The resulting U7snRNA fragments were cloned between the inverted terminal repeats (ITRs) of a self-complementary AAV vector (pscAAV, Addgene plasmid #83279; gift from Mark Kay) for subsequent AAV production. AAV9-pseudotyped vectors were generated by triple co-transfection of HEK293T cells with the pAAV-U7 plasmid, pXX6 plasmid encoding adenoviral helper functions, and pAAV2/9pITRCO2 plasmid containing the *AAV9* rep and cap genes. Seventy-two hours post-transfection, scAAV9-U7-Ex51 vector particles were purified from cell lysates using iodixanol gradient ultracentrifugation. Viral genome titers were determined by quantitative real-time PCR as previously described^39^.

### Administration mode, experimental groups and tissue collection

#### Systemic administration

Intravenous (IV) injections were performed in male *mdx52* mice and wild-type (WT) littermates on postnatal day 2 (P2). Pups were anesthetized by hypothermia induced through 2–3 min exposure to ice. A total volume of 30 µL was injected into the temporal vein. Mice received either ASO-51 (1.2 mg per animal; *n* = 11) or scAAV9-U7-Ex51 (2E+12 vector genomes [vg] per animal; *n* = 11). Control groups received 30 µL of phosphate-buffered saline (PBS) (*mdx52*, *n* = 3; WT, *n* = 3).

#### Intracerebroventricular administration

Intracerebroventricular (ICV) injections were performed in male *mdx52* mice and WT littermates on postnatal day 1 (P1) under hypothermia anaesthesia induced by 2–3 min exposure to ice. ASO-51 or scAAV9-U7-Ex51 was bilaterally injected into the lateral ventricles using surface landmarks visible through the skin at P1 (approximately 0.8–1 mm lateral to the sagittal suture, midway between bregma and lambda, and 3 mm below the skin surface). Each ventricle received 2 µL at an infusion rate of 2 µL/min, corresponding to a total injection volume of 4 µL per animal. This resulted in the administration of 160 µg of ASO-51 (*n* = 17) or 7E+11 vg of scAAV9-U7-Ex51 (*n* = 12). Control animals received 4 µL of sterile PBS (*mdx52*, *n* = 3; WT, *n* = 3).

#### Animal housing

Pups were maintained with their dam and littermates until weaning. Following weaning, male mice were housed in groups of two to five per cage under a 12 h light/12 h dark cycle (lights on at 07:00 h), with ad libitum access to food and water.

#### Tissue collection

At the experimental endpoint, animals were euthanized and tissues were collected for molecular analyses. Brains were dissected to isolate the hippocampus (HIP), cortex (CX), cerebellum (CBL), and cervical spinal cord (SC). Peripheral tissues, including the tibialis anterior (TA), diaphragm (DIA), and heart, were also harvested to evaluate the peripheral biodistribution of ASO-51 and scAAV9-U7-Ex51 following central nervous system administration. All tissue samples were snap-frozen in liquid nitrogen and stored until further processing for biodistribution studies, RT-qPCR, and Western blot analyses.

### RNA analysis

Total RNA was isolated from dissected brain structures, muscles, or cultured cells using TRIzol reagent according to the manufacturer’s instructions (Thermo Fisher Scientific, Rockford, IL, USA). For visualization of exon-skipping efficacy on agarose gels, 1 μg of total RNA was subjected to RT-PCR using the LunaScript RT SuperMix Kit (New England Biolabs, Evry, France) in a 20 μL reaction. Reverse transcription was performed at 55 °C for 10 min. Subsequently, 1.5 μL of cDNA was used for PCR amplification with GoTaq G2 Colorless Master Mix (Promega, Madison, WI, USA) in a 25 μL reaction using the primers Ex-m50F (5′-AGGAAGTTAGAAGATCTGAGG-3′) and Ex-m55R (5′-GGAACTGCTGCAGTAATCTATGA-3′). PCR was performed for 32 cycles of 95 °C for 30 s, 58 °C for 1 min, and 72 °C for 1 min. PCR products were separated on 1.5% agarose gels and quantified using Image Lab Software version 6.1 (Bio-Rad, Marnes-la-Coquette, France). Exon 51 skipping was calculated as the ratio of the skipped band to the native band and expressed as a percentage. Exon 51 skipping was also quantified by TaqMan quantitative PCR as previously described, using TaqMan assays designed against the exon 50–51 junction (assay Mm.PT.58.41685801: Forward: 5′-CAAAGCAGCCTGACCGT-3′; Reverse: 5′-TGACAGTTTCCTTAGTAACCACAG-3′; Probe: 5′-TGGACTGAGCACTACTGGAGCCT-3′) and the exon 50–53 junction (Forward: 5′-GCACTACTGGAGCCTTTGAA-3′; Reverse: 5′-CTTCCAGCCATTGTGTTGAATC-3′; Probe: 5′-ACAGCTGCAGAACAGGAGACAACA-3′) (Integrated DNA Technologies). A total of 150 ng of cDNA was used per reaction, and all assays were performed in triplicate. Reactions were run under fast cycling conditions on a Bio-Rad CFX384 Touch Real-Time PCR Detection System, and data were analyzed using the absolute copy number method. For each sample, copy numbers of the skipped product (exon 50–53) and unskipped product (exon 50–51) were determined using the standards Ex49-54Delta52 and Ex49-54Delta51+52 (gBlocks gene fragments from Integrated DNA Technologies). Exon 51 skipping was expressed as a percentage of total dystrophin, calculated as the sum of exon 50–51 and exon 50–53 copy numbers.

### Protein Analysis

Protein extracts were prepared from brain tissues using RIPA lysis and extraction buffers (Thermo Fisher Scientific), supplemented with SDS to a final concentration of 5% (Bio-Rad, France). Protein concentrations were determined using the BCA Protein Assay Kit (Thermo Fisher Scientific).

For conventional Western blot analyses, samples were denatured at 100 °C for 3 min, and 25 µg of total protein per sample was loaded onto NuPAGE 3–8% Tris-acetate gels (Invitrogen) according to the manufacturer’s instructions. Dystrophin expression was detected using a rabbit monoclonal anti-dystrophin antibody (ab154168, Abcam, France), while vinculin was used as a loading control and detected with the mouse monoclonal hVin-1 antibody (V9131, Sigma-Aldrich, USA). Membranes were incubated with IRDye 680CW goat anti-rabbit and IRDye 800CW goat anti-mouse secondary antibodies (LI-COR Biosciences, Germany). Fluorescent signals were acquired using an Odyssey CLx imaging system (LI-COR), and quantification was performed with Image Studio software (LI-COR). Dystrophin expression levels were normalized to vinculin.

For samples from the systemic scAAV9-U7-Ex51 study, dystrophin expression was assessed using an automated capillary-based Western blot system (Jess™, ProteinSimple, Bio-Techne, USA), which provides automated protein separation and immunodetection while reducing sample consumption and operator-dependent variability. Protein extracts were prepared as described above and analyzed using the 66–440 kDa separation module. Samples, primary antibodies, and detection reagents were loaded into the assay plate and processed automatically by the instrument according to the manufacturer’s instructions. The same anti-dystrophin (ab154168, Abcam) and anti-vinculin (V9131, Sigma-Aldrich) antibodies were used. Chemiluminescent signals for dystrophin and near-infrared (NIR) fluorescence signals for vinculin were analyzed using Compass for Simple Western software (ProteinSimple), and dystrophin expression levels was normalized to vinculin levels.

For both conventional Western blot and Jess analyses, standard curves were generated for each brain region by mixing lysates from wild-type (WT) and *mdx52* control samples to obtain defined dystrophin levels corresponding to 0%, 2.5%, 5%, 10%, 20%, 30%, 50% and 100% of WT expression.

### Statistical analysis

The data are presented as means ± standard error of the mean (SEM). The effects of timepoints and brain regions were analyzed using the nonparametric two-way ANOVA followed by Tukey’s multiple comparisons test. A p-value of <0.05 was considered statistically significant.

## Supporting information

Supplemental informations

## DATA AVAILABILITY STATEMENT

The data that support the findings of this study are available from the corresponding authors upon reasonable request.

## SUPPLEMENTAL INFORMATION

Supplemental information includes Figures S1 to S4:

Figure S1. Molecular outcomes in the spinal cord following neonatal ASO-51 or scAAV9-U7-Ex51 administration.

Figure S2. Representative western blot and Jess images used for dystrophin quantification following neonatal ASO-51 or scAAV9-U7-Ex51 administration.

Figure S3. Molecular outcomes in peripheral tissues following ICV administration of ASO-51.

Figure S4. Molecular outcomes in peripheral tissues following ICV administration of scAAV9-U7-Ex51.

## ACKNOWLEDGMENTS

This work was partly funded by the European Union’s Horizon 2020 research and innovation program “Brain Involvement iN Dystrophinopathies” under grant agreement No 847826. It was also supported by Agence Nationale pour la recherche (grant BRAIN-DYS ANR-24-CE18-1633), Institut National de la santé et la recherche médicale (INSERM), Université Paris-Saclay (France) and Paris Ile-de-France Region. We thank Dr. Katsuki Motoya, Prof. Sasaoka Toshikuni (Department of Comparative & Experimental Medicine/ Brain Research Institute; Niigata University, Japan), Dr. Jun Tanihata and Dr. Shin’ichi Takeda (National Center of Neurology and Psychiatry, Tokyo, Japan) for providing the *mdx52* mouse breeders. We gratefully acknowledge the Pelvipharm team for the management, housing, and care of the mouse colonies used throughout this study.

## AUTHOR CONTRIBUTIONS

Project design, funding acquisition and resources. A.G. and C.V.; Design of experiments. A.G. and O.V.; Production of AAV vectors. O.V. and X.P; Experimental work: O.V. M.D. C.N. A.S. X.P. and O.L.C.; Data analysis. A.G. and O.V.; Original draft preparation. O.V. A.G. and M.D.; All authors have read and agreed to the published version of the manuscript.

