## Supplemental informations for "Neonatal Delivery of Exon Skipping Therapeutics Maximizes Brain Dystrophin Rescue in the *Mdx52* Mouse Model of Duchenne Muscular Dystrophy"

### Supplemental Material Vacca et al

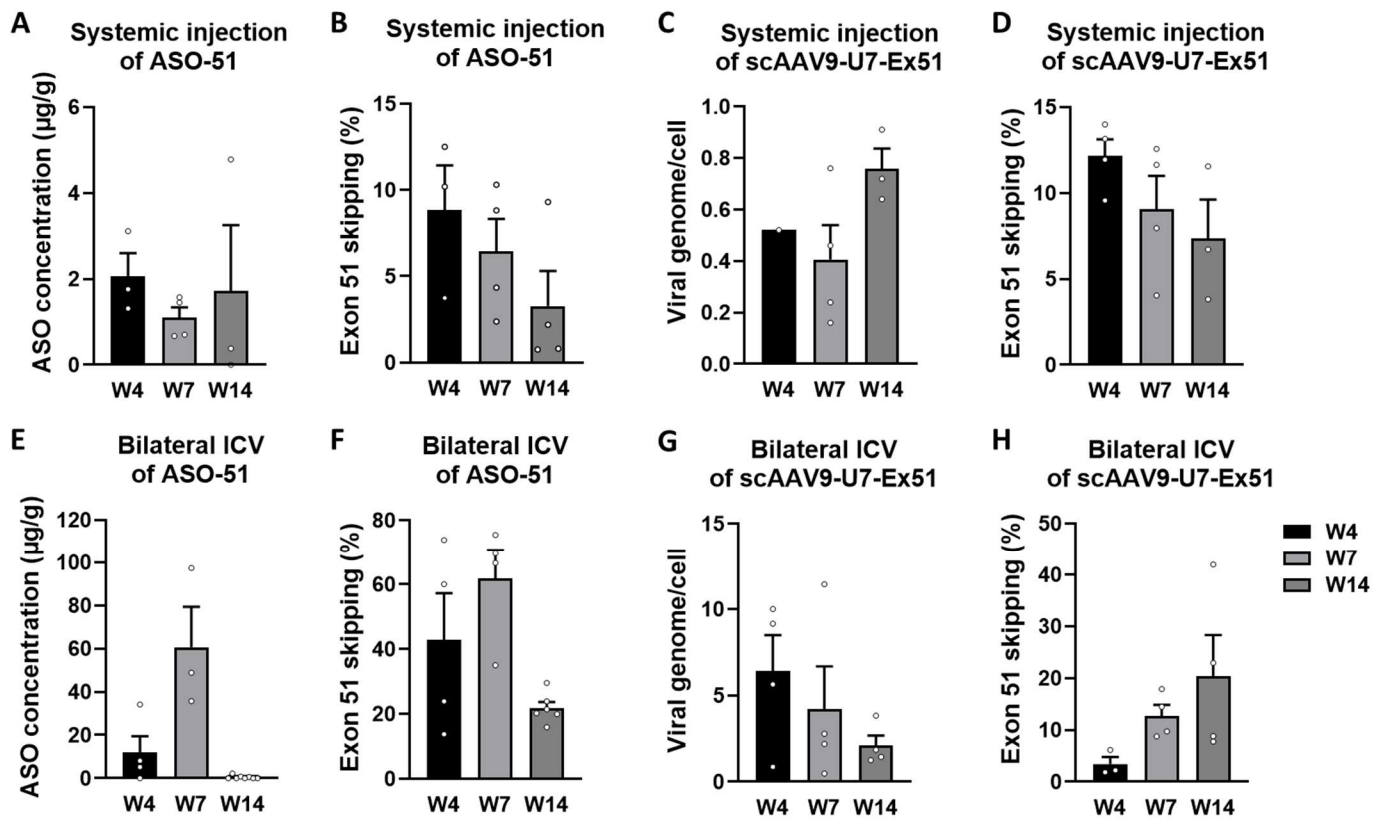

**Supplementary Figure S1. Molecular outcomes in the spinal cord following neonatal ASO-51 or scAAV9-U7-Ex51 administration.** Spinal cord samples were collected at weeks 4 (W4), 7 (W7), and 14 (W14) after neonatal treatment. (A, B) ASO concentration (A) and exon 51 skipping levels (B) following systemic administration of ASO-51. (C, D) Viral genome copy number per cell (C) and exon 51 skipping levels (D) following systemic administration of scAAV9-U7-Ex51. (E, F) ASO concentration (E) and exon 51 skipping levels (F) following bilateral intracerebroventricular (ICV) administration of ASO-51. (G, H) Viral genome copy number per cell (G) and exon 51 skipping levels (H) following bilateral ICV administration of scAAV9-U7-Ex51. Bars represent mean ± SEM, and individual animals are shown as dots. Spinal cord data are shown separately as a supplementary figure for completeness.

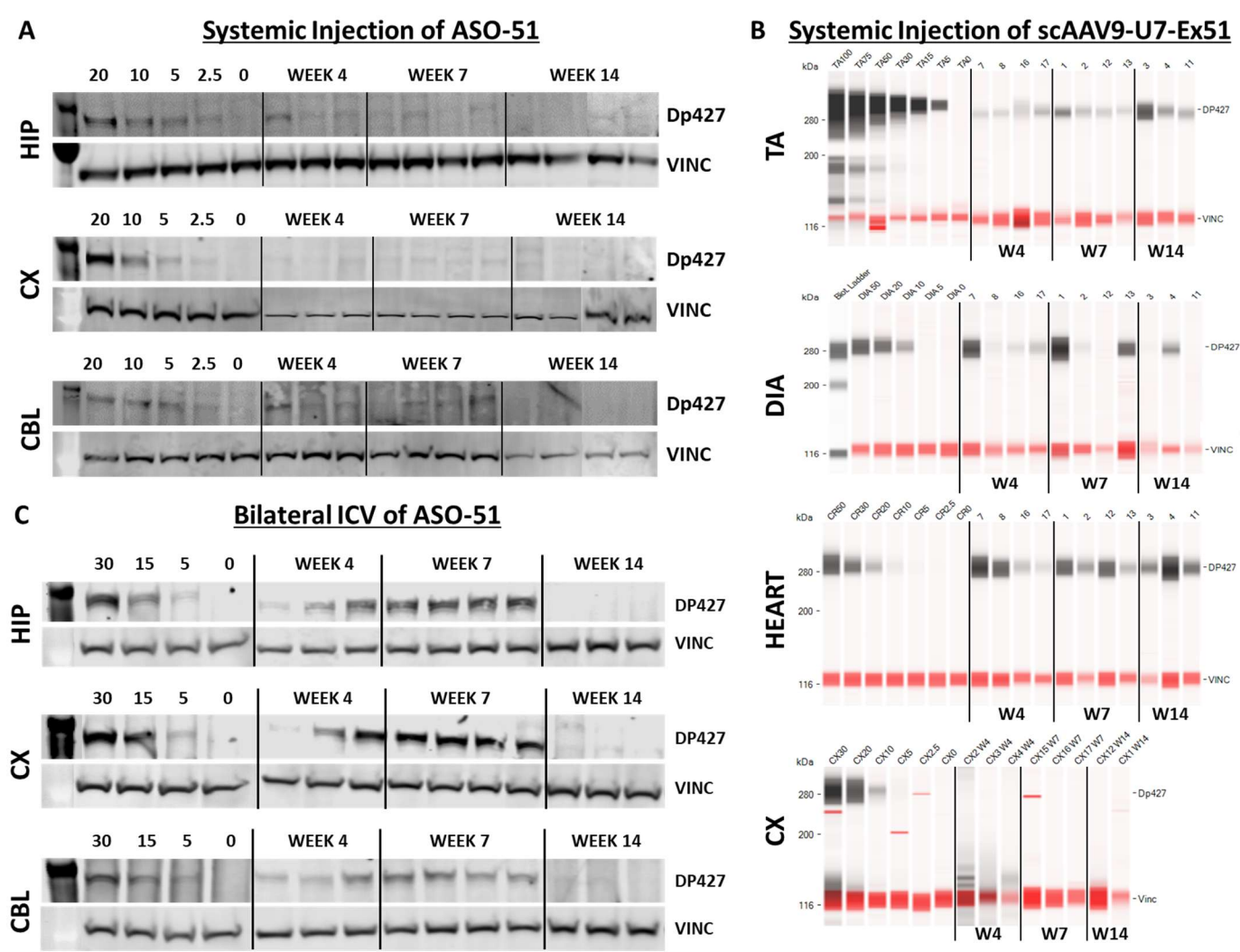

**Supplementary Figure S2. Representative western blot and Jess images used for dystrophin quantification following neonatal ASO-51 or scAAV9-U7-Ex51 administration.** (A) Representative western blot images showing Dp427 expression in the hippocampus (HIP), cortex (CX), and cerebellum (CBL) following systemic neonatal administration of ASO-51. Vinculin (VINC) was used as the loading control. Lanes on the left correspond to the dystrophin standard curve (20, 10, 5, 2.5 and 0% of wild-type protein), followed by samples collected at weeks 4, 7, and 14 after injection. (B) Representative capillary western immunoassay (Jess) images showing Dp427 expression in tibialis anterior (TA), diaphragm (DIA), heart and cortex (CX) following systemic neonatal administration of scAAV9-U7-Ex51. The cortex is shown as the representative CNS tissue, as it displayed the highest exon-skipping levels following treatment. Despite this, no restored Dp427 protein was detectable. Vinculin (VINC) signals in red were used as the loading control. The standard curve (100, 50, 30, 20, 10, 5 and 0% of wild-type protein) is shown on the left, followed by samples were collected at weeks 4, 7, and 14 after injection. (C) Representative western blot images showing Dp427 expression in the hippocampus (HIP), cortex (CX), and cerebellum (CBL) following bilateral neonatal intracerebroventricular (ICV) administration of ASO-51. Vinculin (VINC) was used as the loading control. The standard curve (30, 15, 5 and 0% of wild-type protein) is shown on the left, followed by samples collected at weeks 4, 7, and 14 after injection. All images are representative of those used for densitometric analyses presented in the main figures.

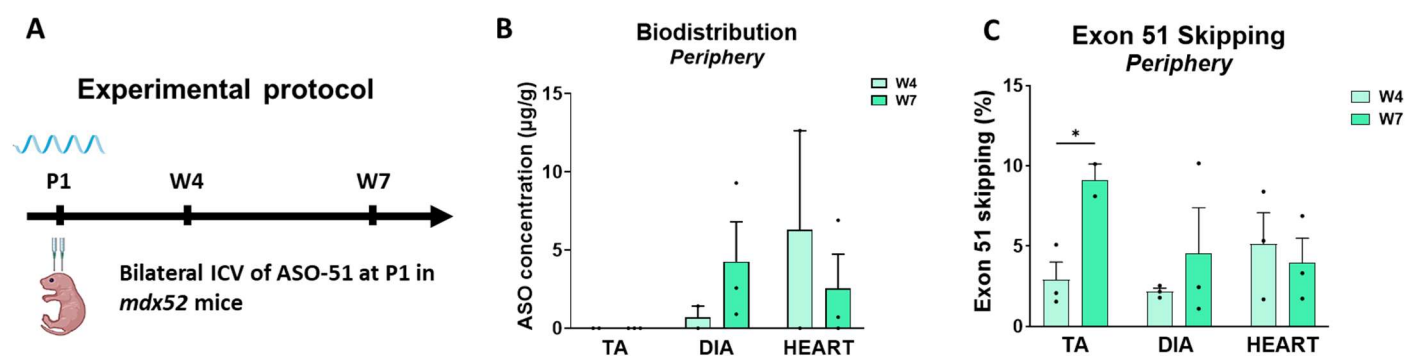

**Supplementary Figure S3. Molecular outcomes in peripheral tissues following ICV administration of ASO-51.**

(A) Experimental timeline. *Mdx52* mice received a single bilateral intracerebroventricular (ICV) injection of ASO-Ex51 at postnatal day 1 (P1). (B) ASO-51 biodistribution in peripheral tissues, including tibialis anterior (TA), diaphragm (DIA), and heart at W4 and W7. (C) Exon 51 skipping quantification by RT-qPCR in tibialis anterior (TA), diaphragm (DIA), and heart at W4 and W7. Data are presented as mean ± SEM with individual data points shown ($n=3$ per time point). Statistical analysis was performed using a two-way ANOVA followed by Tukey's multiple comparisons test. \*$p < 0.05$.

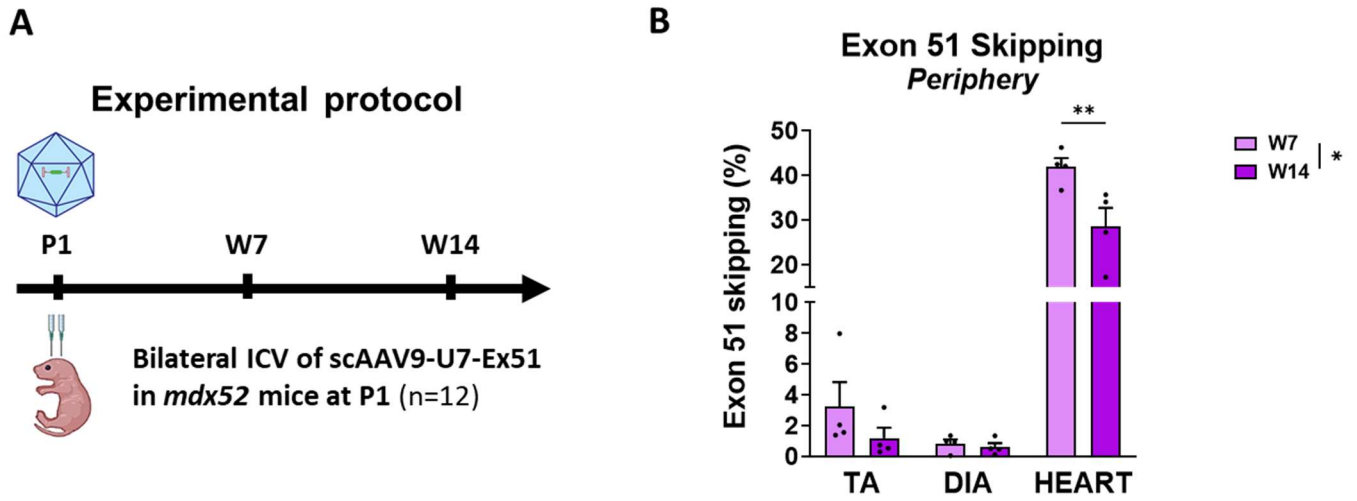

**Supplementary Figure S4. Molecular outcomes in peripheral tissues following ICV administration of scAAV9-U7-Ex51.** (A) Experimental timeline. *Mdx52* mice received a single bilateral intracerebroventricular (ICV) injection of scAAV9-U7-Ex51 at postnatal day 1 (P1). (B) Exon 51 skipping quantification by RT-qPCR in tibialis anterior (TA), diaphragm (DIA), and heart at W7 and W14. Data are presented as mean ± SEM with individual data points shown (n=4 per time point). Statistical analysis was performed using a two-way ANOVA followed by Tukey's multiple comparisons test. \*$p < 0.05$, \*\*$p < 0.01$.
